# Hyperactive end joining repair mediates resistance to DNA damaging therapy in p53-deficient cells

**DOI:** 10.1101/2020.04.01.021253

**Authors:** Rashmi J. Kumar, Hui Xiao Chao, Victoria R. Roberts, Aurora R. Sullivan, Sonam J. Shah, Dennis A. Simpson, Wanjuan Feng, Anne-Sophie Wozny, Sunil Kumar, Jeremy E. Purvis, Gaorav P. Gupta

**Author notes:** **Correspondence:** Gaorav P. Gupta, MD PhD, Assistant Professor, Department of Radiation Oncology, Department of Biochemistry and Biophysics, University of North Carolina at Chapel Hill, Chapel Hill, NC 27599.

## Abstract

*TP53* mutations in cancer are associated with poor patient outcomes and resistance to DNA damaging therapies^1–3^. However, the mechanisms underlying treatment resistance in p53-deficient cells remain poorly characterized. Here, we show that p53-deficient cells exhibit hyperactive repair of therapy-induced DNA double strand breaks (DSBs), which is suppressed by inhibition of DNA-dependent protein kinase (DNA-PK). Single-cell analyses of DSB repair kinetics and cell cycle state transitions reveal an essential role for DNA-PK in suppressing S phase DNA damage and mitotic catastrophe in p53-deficient cells. Yet, a subset of p53-deficient cells exhibit intrinsic resistance to therapeutic DSBs due to a repair pathway that is not sensitive to DNA-PK inhibition. We show that p53 deficiency induces overexpression of DNA Polymerase Theta (Pol θ), which mediates an alternative end-joining repair pathway that becomes hyperactivated by DNA-PK inhibition^4^. Combined inhibition of DNA-PK and Pol θ restores therapeutic DNA damage sensitivity in p53-deficient cells. Thus, our study identifies two targetable DSB end joining pathways that can be suppressed as a strategy to overcome resistance to DNA-damaging therapies in p53-deficient cancers.

## Introduction

*TP53* is the most commonly mutated tumor suppressor gene^5^. p53 mediates pleiotropic tumor suppressive effects through regulation of cell cycle arrest, apoptosis, and cellular metabolism in response to cellular stress^6,7^. Beyond its role as a tumor suppressor, loss of functional p53 is associated with poor prognostic outcomes across many different cancer types^1,8–10^. There is both clinical and preclinical evidence that p53-deficient cancers exhibit resistance to a variety of DNA damaging therapies^2,3,11–14^.

The mechanisms for therapeutic resistance in p53-deficient cells remains poorly characterized. Past work has suggested a role for loss of p53-mediated apoptosis^12,15^. However, the response of epithelial cancer cells to DNA damaging therapy is often determined by the efficiency of inducing senescence or mitotic catastrophe, rather than apoptosis^16,17^. p53 is also a transcription factor that responds to DNA double strand breaks (DSBs) to determine cellular fate^7,18^. Recent insights have revealed the importance of p53-signaling waves in regulation of cellular fate decisions of quiescence versus cell cycle re-entry after DNA damage^19,20^. However, the mechanisms that determine such cell fate decisions upon DNA damage induction in p53-mutant epithelial cells have not been established, and may lead to novel strategies to restore treatment sensitivity.

In this study, we investigate altered DNA repair mechanisms in p53 deficiency as a major contributor to resistance to DNA damaging therapies. We find that p53-deficient cells exhibit hyperactive repair and accelerated resolution of DNA damage foci. Utilizing live-cell imaging, we show that this ability to resolve DNA damage rapidly is partially dependent on DNA-PK, a critical serine/threonine kinase in the non-homologous end joining (NHEJ) pathway^21^. Inhibition of DNA-PK using the small molecule inhibitor NU7441 partially sensitizes p53-deficient cells to DSB inducing agents. We further show that this effect is specifically due to propagation of S phase related damage leading to mitotic catastrophe, highlighting a role for DNA-PK in S phase DNA damage repair that was previously under appreciated. Furthermore, using chromosomal break repair assays we show that in the context of inhibitor treatment, some p53-deficient cells utilize alternative end-joining repair in a compensatory manner to escape cell death. Thus, our work provides critical insight into a clinically-relevant mechanism for why p53-deficient cells are resistant to DNA damaging therapies.

## Results

### p53-deficient cells exhibit radioresistance and accelerated resolution of DNA DSBs

We first established an isogenic cell system to investigate determinants of treatment-induced cell fate in p53-deficient cells. In order to minimize potential contributions of accessory mutations on phenotypes observed in cancer cell line models, we used CRISPR/Cas9 to disrupt *TP53* in the p53-proficient immortalized epithelial cell line model hTert-RPE1 (“RPE1”), which has also been a preferred model for investigating p53-dependent cell fate^18,20,22^. Two independent CRISPR/Cas9-targeted *TP53^−/−^* RPE1 clones were selected for further study after confirming cells were deficient for p53 protein and lacked p53-dependent transcriptional induction of p21 in response to ionizing radiation (IR) (Supplementary Fig. 1a-c).

To assess whether p53 deficiency confers a proliferative advantage when treated with ionizing radiation, we performed a mixed competition assay. We took mCherry labelled RPE1 and mixed them with equal numbers of unlabeled *TP53^−/−^* RPE1 or p53-proficient RPE1 (control) (Fig. 1a). We quantified the relative abundance of the unlabeled cells after to exposure to IR (0 – 6Gy), normalized to untreated samples at each timepoint. RPE1 labeled and unlabeled cells maintained stable representation across time and treatment conditions (Supplementary Fig. 1d). Additionally, p53-deficient cells did not demonstrate a proliferation advantage in the absence of RT. However, treatment with IR at any dose level led to substantial positive selection for p53-deficient cells(Fig. 1b,c). We also observed that p53 deficiency induced resistance to the radiomimetic clastogen, Neocarzinostatin (NCS) by colony forming assay (Supplementary Fig. 1e-i). Thus p53 deficiency in this isogenic model is sufficient to induce radioresistance.

**Figure 1.**
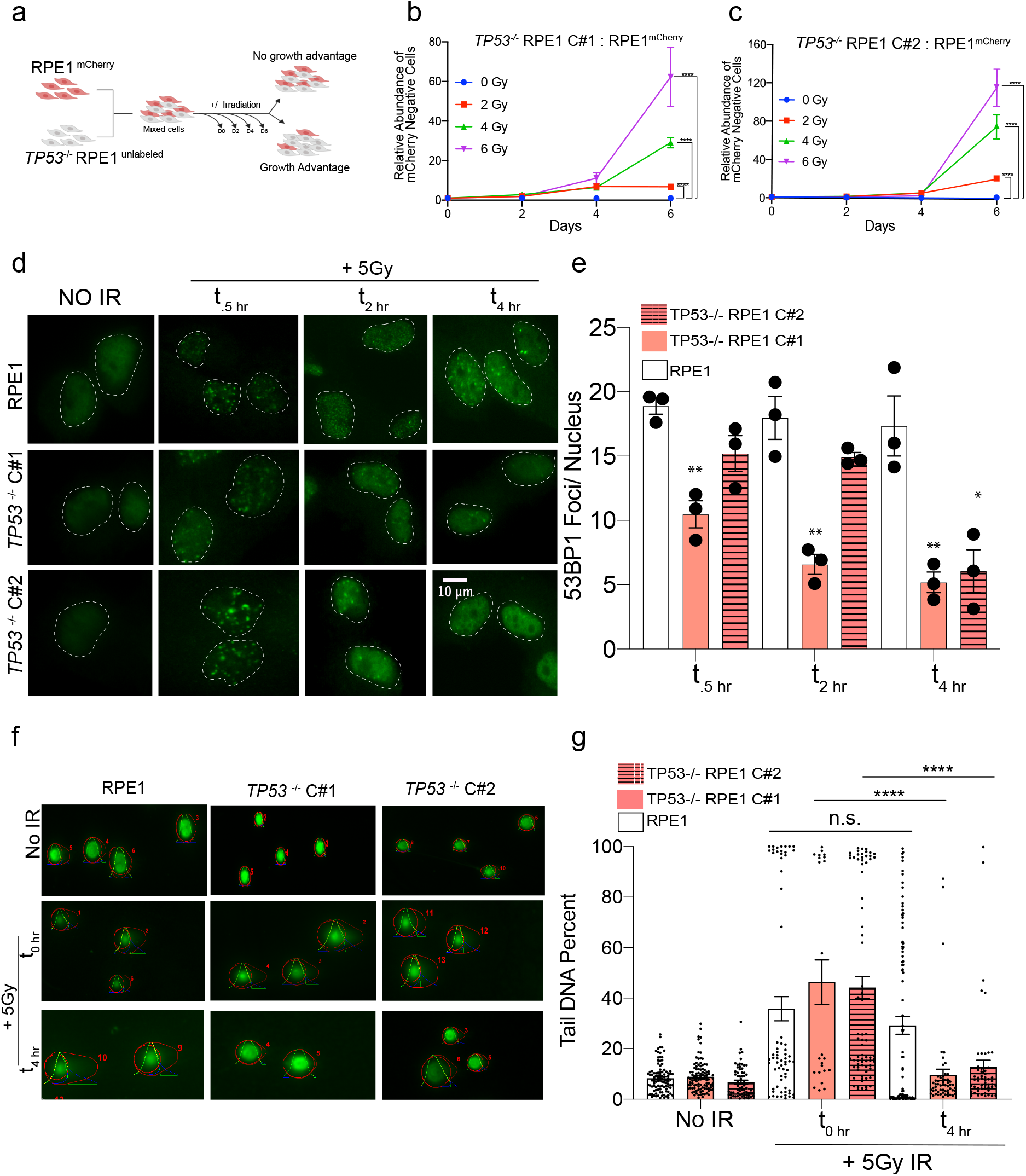
p53-deficient cells exhibit radioresistance and accelerated resolution of DNA DSBs. **a,** Diagram of growth competition assay. mCherry-labelled RPE1 cells were mixed with unlabeled *TP53*^−/−^ RPE1 (1:1), exposed to IR and grown for 6 days. **b,** Relative abundance of unlabeled *TP53* ^−/−^ Clone#1 measured by Intellicyte high-throughput cytometry ± SEM (n=6) is shown, normalized to the untreated (0Gy) cohort at each time point. **c,** Relative abundance of unlabeled *TP53* ^−/−^ Clone#2 ± SEM (n=6) is shown, normalized to the untreated (0Gy) cohort at each time point. **d,** Representative immunofluorescence images of 53BP1 foci in cells with indicated genotypes untreated (no IR) or treated with IR (5Gy) and collected at .5, 2, and 4 h after irradiation. **e,** Quantification of 53BP1 foci. Data shown are mean (n=50 cells per treatment condition) ± SEM (n=3), and are consistent across two independent biological replicates. \**p*<0.05; \*\**p*<0.01; by two-tailed Student’s t-test. **f,** Representative Neutral COMET fluorescence staining for DNA tails in cells with indicated genotypes treated with or without 5Gy IR. For irradiated cells, 2 timepoints are shown: immediately after and 4 hours post IR. COMET tails and heads are denoted by OpenComet software analysis. **g,** Quantification of DNA DSBs via Neutral COMET assay reported as tail DNA percent at 0 and 4 hours post IR in RPE1 and two *TP53*^−/−^ RPE1 cell lines. Data shown are mean (n= 50-150 cells per treatment condition) ± SEM, and are consistent across three independent biological replicates. \**p*<0.05; \*\**p*<0.01; \*\*\*\**p*<0.0001 by two-tailed Student’s t-test.

Unrepaired DSBs can suppress proliferation through the engagement of DNA damage-induced cell cycle checkpoints. We examined kinetics of DSB repair by performing immunofluorescence for 53BP1 and γH2AX after treatment of p53 WT and *TP53^−/−^* cells with 5Gy IR (Fig. 1d,e and Supplementary Fig. 1j,k). We observed a reduction in the number of 53BP1 damage foci in *TP53^−^*^/-^ cells as early as 30 minutes after treatment, that became even more pronounced by 4 hours post-treatment (Fig. 1d,e). Similar patterns of reduced foci formation were also apparent with γH2AX staining at early timepoints (Supplementary Fig. 1j,k). Quantification of IR-induced DSBs by neutral COMET assay revealed an equivalent DSB burden induced immediately after 5Gy IR, irrespective of p53 status (Fig. 1f,g). However, by 4 hours post-treatment, tail DNA percent was significantly reduced in the *TP53^−/−^* cells while remaining elevated in p53-proficient RPE1 (Fig. 1f,g). Thus, p53 deficiency is sufficient to induce radioresistance and accelerated DSB repair in an isogenic model.

### Inhibition of DNA-PK restores DNA damage foci formation in p53-deficient cells

To directly assess the relationship between DSB repair kinetics, cell cycle status, and cell fate at the single cell level, we established a live cell imaging platform (Fig. 2a). RPE1 cells were dually labeled with PCNA-mCherry (to monitor cell cycle state transitions) and 53BP1-mVenus (to monitor DSB foci kinetics) (Fig. 2b)^23,24^. These dual labeled cells were treated with scrambled siRNA (si-Control) or siRNA targeting *TP53* (si-*TP53*), the latter of which resulted in >90% knockdown of *TP53* transcript and elimination of p53-dependent *CDKN1A* transcription in response to IR (Fig. 2c). 48 hours after siRNA treatment, RPE1 cells were imaged for a total of 72 hours every 10 minutes, and 18 hours into imaging, the DSB inducing agent was added (Fig. 2a). To minimize time from radiation exposure to image capture and to induce equivalent DSBs in each population of cells, we utilized 100 ng/ml of Neocarzinostatin (NCS), a well-known radio-mimetic. NCS has been previously utilized in studies evaluating DNA DSB repair in conjunction with live-cell imaging and has been shown to induce peak DSBs within 10 minutes of drug addition^25,26^. This experimental design allowed us to determine the cell cycle status of each cell within the asynchronous cell population at the time of NCS exposure. After NCS treatment, single-cell analyses for DSB repair foci kinetics and cell cycle outcomes were performed. As anticipated from Fig. 1b,c, analysis of global proliferation by live-cell imaging revealed significantly greater proliferation of p53-deficient RPE1 cells relative to controls after NCS treatment (Supplementary Fig. 2a,b).

**Figure 2.**
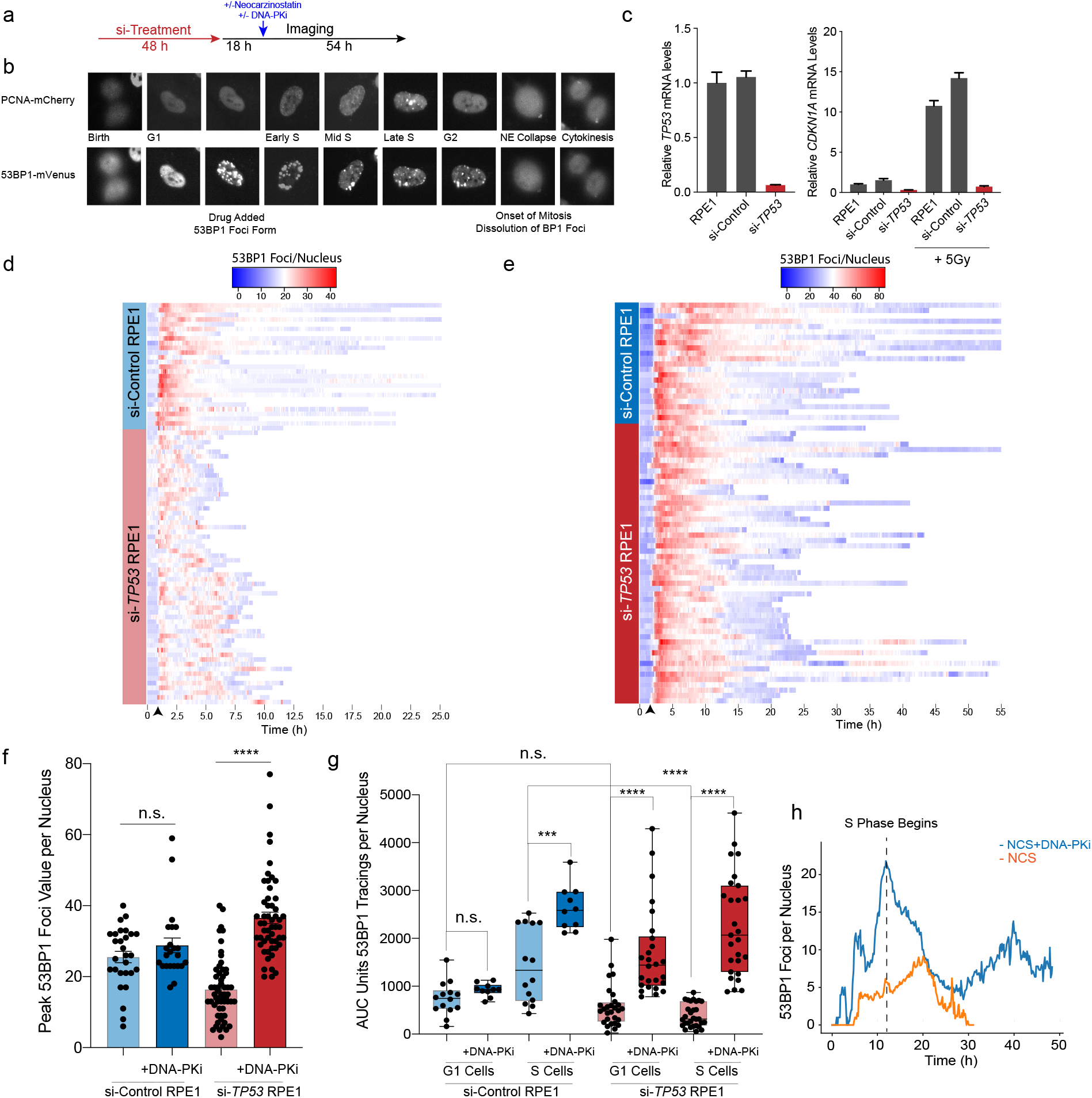
Inhibition of DNA-PK restores DNA damage foci formation in p53-deficient cells. **a,** Live cell imaging procedure. Cells transfected with 10 nM si-control or si-*TP53* for 48 h prior to imaging. 18 h into imaging, cells are treated with NCS (100 nM), DNA-PKi (.5 uM) or both and imaged for 72 total hours. **b,** RPE1 cell expressing the PCNA-mCherry and 53BP1-mVenus reporters. Cell cycle phases delineated by PCNA foci and DNA DSBs are marked by 53BP1 foci. **c,** RT-qPCR for *TP53* mRNA levels (left) and *CDKN1A* mRNA levels (right) in si-control treated vs. si-*TP53* treated cells. To induce *CDKN1A* expression, cells irradiated at 5Gy and mRNA harvested 3 hrs post IR. **d,** Heatmap of 53BP1 foci tracings for single cells tracked from birth to mitosis or end of imaging. For si-control (n = 30 cells) and si-*TP53* treated RPE1 (n = 60 cells) treated with NCS 100 ng/ml. For visualization, cells are aligned to 10 frames prior to drug addition (black arrow). **e,** Heatmap of 53BP1 foci tracings for si-control (n = 25 cells) and si-*TP53* treated RPE1 cells (n = 55 cells) treated with 100 ng/ml NCS + 0.5 uM DNA-PKi. **f,** Peak 53BP1 foci counts for cells treated with 100 ng/ml NCS or NCS+0.5 uM DNA-PKi. Significance determined using two-tailed t-test. **g,** Area under the curve (AUC) analysis of 53BP1 burden showing integral DNA damage for cells treated with NCS vs. NCS and DNA-PKi. Cells are segregated into two groups: cells exposed to drug in G1 vs. S phase (n = 25-30 G1or S cells for si-*TP53* cohort, n = 10-15 G1 or S cells for si-control cohort). Significance determined by two-tailed t-test. \*\*\*\**p*<0.0001, \*\*\**p*<0.001, n.s. = non-significant. **h,** 53BP1 foci burden in G1 vs. S phase p53-deficient RPE1 upon exposure to NCS and DNA-PKi. Dashed line = S phase onset, blue line = mean 53BP1 foci burden for all cells in G1 with NCS and DNA-PKi addition, orange line = mean foci value for cells in G1 with NCS treatment alone, (n = 30 cells for each condition).

To analyze DSB repair kinetics in cells exposed to NCS, we tracked and quantified 53BP1 foci in single cells and plotted heatmaps of damage foci burden over time from cell birth to mitosis (Fig. 2d). Our results indicate that cells with functional p53 sustain high levels of damage foci in a prolonged manner after NCS exposure. In contrast, p53-deficient cells developed a lower peak burden of 53BP1 foci after NCS treatment, with accelerated resolution of damage foci to baseline levels (Fig. 2d,f). Given the rapidity with which 53BP1 foci were being resolved, we hypothesized that hyperactive NHEJ may be contributing. We thus performed the same experiment in the presence of an inhibitor of DNA-dependent Protein Kinase (DNA-PKi, NU7441 0.5μM), which targets the central kinase in the NHEJ pathway^27–29^. Strikingly, DNA-PKi qualitatively abolished the difference in 53BP1 kinetics after NCS treatment between p53-deficient and proficient cells (Fig. 2e). To quantitatively assess the magnitude in damage burden, we calculated peak maximum 53BP1 foci values for each cell represented in the heatmap (Fig. 2f). Consistent with the heatmap representation, the median peak foci count after NCS treatment was 40% lower in si-*TP53* treated cells relative to controls (Fig. 2f, p<0.0001). Notably, DNA-PKi treatment resulted in a >2-fold increase in peak 53BP1 foci levels in the p53-deficient cells, whereas there was no comparable effect in control cells (Fig. 2f). These results indicate that DNA-PK activity is required for accelerated resolution of clastogen-induced DNA damage foci in p53-deficient cells.

Following this analysis, we studied the effects of DNA-PKi in different phases of the cell cycle during drug exposure. We used PCNA live-cell imaging to resolve cell cycle phase transitions in cells tracked for 53BP1 foci kinetics. We performed area under the curve (AUC) analyses in single cells to estimate total DNA damage burden during G1 and S phase after NCS exposure (Fig. 2g). This analysis revealed that the diminished 53BP1 foci burden observed in p53-deficient cells was most pronounced during S phase relative to control cells (Fig. 2g). DNA-PKi treatment significantly increased S phase 53BP1 burden in both si-Control and si-*TP53* treated RPE1 cells (Fig. 2g). While si-*TP53* treated cells in G1 were also affected to a lesser degree, we were curious to examine if the effect was in part due to loss of the p53-dependent G1/S checkpoint resulting in propagation of unrepaired DNA damage into S phase. Indeed, we found that DNA-PKi induced a drastic increase in 53BP1 foci as p53-deficient cells transitioned from G1 to S phase, which subsequently diminished over time (Fig. 2h, p<0.00001 at t = start of S phase). Thus, DNA-PK is required for hyperactive resolution of clastogen-induced DSB foci in p53-deficient cells, and most prominently during S phase.

### Checkpoint responses halt p53-proficient cells upon exposure to NCS while p53-deficient cells continue to cell cycle despite NCS exposure

To investigate the association between DNA damage and activation of cell cycle checkpoints, we quantified cell cycle phase durations for all treatment conditions (Fig. 3a,b). p53-proficient G1 cells exposed to NCS induced a significant prolongation of G1, indicative of G1/S checkpoint activation, with a substantial proportion of cells remaining arrested for the duration of imaging (Fig. 3c and Supplementary Fig. 3a-c). Similarly, cells exposed to NCS in S phase exhibited a G2-M checkpoint (Fig. 3d). p53-deficient cells exhibited no prolongation of G1 duration after NCS, consistent with the notion that G1/S checkpoint activation is p53-dependent (Fig. 3e)^30,31^. DNA-PK inhibition did not alter G1 duration in either p53-proficient or p53-deficient cells (Fig. 3c,e). In contrast, DNA-PKi increased the duration of G2-M checkpoints irrespective of p53 status (Fig. 3d,f). These observations suggest that increased levels of S phase DNA damage induced by DNA-PKi and NCS treatment (see Fig. 2g,h) result in activation of a G2/M checkpoint that is, at least partially, p53-independent. However, the duration of G2/M checkpoint activation differed by p53 status. While p53-proficient cells frequently remained arrested for the entire duration of imaging (open circles, Fig. 3c,d), p53-deficient cells experienced a more transient prolongation of G2 duration followed by progression into mitosis (Fig. 3e,f).

**Figure 3.**
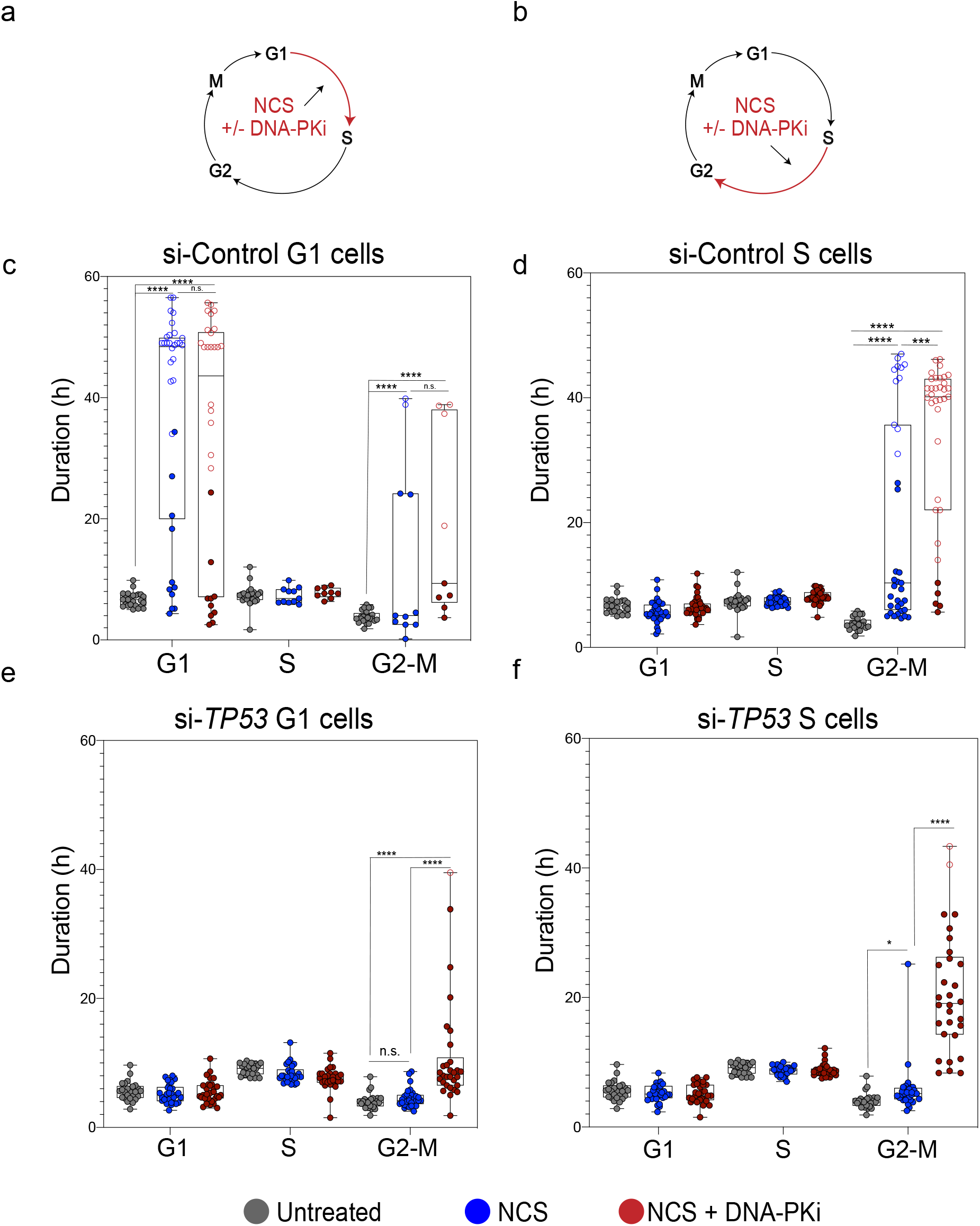
Checkpoint responses halt p53-proficient cells upon exposure to NCS while p53-deficient cells continue to cell cycle despite NCS exposure. **a,** Schematic depicting NCS treatment (50 ng/ml + 100 ng/ml for si-control and 100 ng/ml for si-*TP53* RPE1) and/or NCS + 0.5 uM DNA-PKi treatment, and phase of the cell cycle cells are exposed to drug (G1). **b,** Schematic of drug treatment for S phase cells. **c,** Distribution of cell cycle phase lengths, each colored dot is an individual cell with untreated cells (no NCS) shown in black, NCS treated cells shown in blue, and NCS+ 0.5uM DNA-PKi treated cells shown in red for si-control RPE1 in G1 phase. n = 20 untreated and n = 30 treated cells (for each treatment cohort). Statistical significance was determined by comparing untreated and treated groups at each phase. \*\*\*\**p*<0.0001, n.s. = non-significant. Open circles indicate arrested cells that did not enter the subsequent phase of cell cycle for remainder of imaging. **d,** Distribution of cell cycle phase lengths for si-control treated RPE1 in S phase, \*\*\**p*<0.001, \*\*\*\**p*<0.0001, n.s. = non-significant as evaluated by two-tailed t-test. **e,** Distribution of cell cycle phase lengths for si-*TP53* treated RPE1 in G1 phase, \*\*\*\**p*<0.0001, n.s. = non-significant as evaluated by two-tailed t-test. **f,** Distribution of cell cycle phase lengths for si-*TP53* treated RPE1 in S phase, \*\*\*\**p*<0.0001, n.s. = non-significant as evaluated by two-tailed t-test.

### Inhibition of DNA-PK induces catastrophic mitoses in p53-deficient cells

We next used a heatmap representation to track the fate of individual cells from birth until mitosis (Fig. 4a,b, top panels). Red bars indicate a mitotic catastrophe or apoptosis event (Supplementary Fig. 4a,b). The median cell cycle time for both untreated p53-proficient and p53-deficient cells was approximately 22-24 hours. NCS treatment is indicated as a dashed line at the 18 hour timepoint. Individual cells are ordered according to cell cycle phase at the time of NCS treatment (G1 versus S) and eventual cell fate (viable, G1 arrest, G2 arrest, or mitotic catastrophe/apoptosis). The majority (70%) of p53-proficient (si-Control) G1 cells exposed to NCS activated a G1 checkpoint that was maintained for the remainder of imaging (Fig. 4a). 26% of these cells underwent G2 arrest or mitotic catastrophe, whereas only 3% retained their proliferative capacity (Fig. 4a). Control cells exposed to NCS in S phase exhibited more diverse cell fates: 40% G2 arrest, 17% mitotic catastrophe, and 43% that retained proliferative capacity. These observations, made using single-cell tracking of asynchronous cell populations, are consistent with observations of intrinsic radioresistance of S phase cells using cell synchronization methods^32^. In contrast, the majority of p53-deficient (i.e., si-*TP53* treated) cells in G1 or S at the time of NCS treatment remained viable without perceptible engagement of any cell cycle checkpoints (Fig. 4b, 80% and 87%, respectively). Consistent with prior 53BP1 analyses, S phase cells are most sensitized to DNA-PKi as the addition of the inhibitor increased G2 arrest frequency in control cells (40% to 91%), and increased mitotic catastrophe in p53-deficient cells (13% to 47%, Fig. 4a,b). In total, the percentage of viable p53-deficient cells after NCS decreased from 87% to 47% when treated in S phase with DNA-PK inhibition (p<0.0001, Fisher’s exact test).

**Figure 4.**
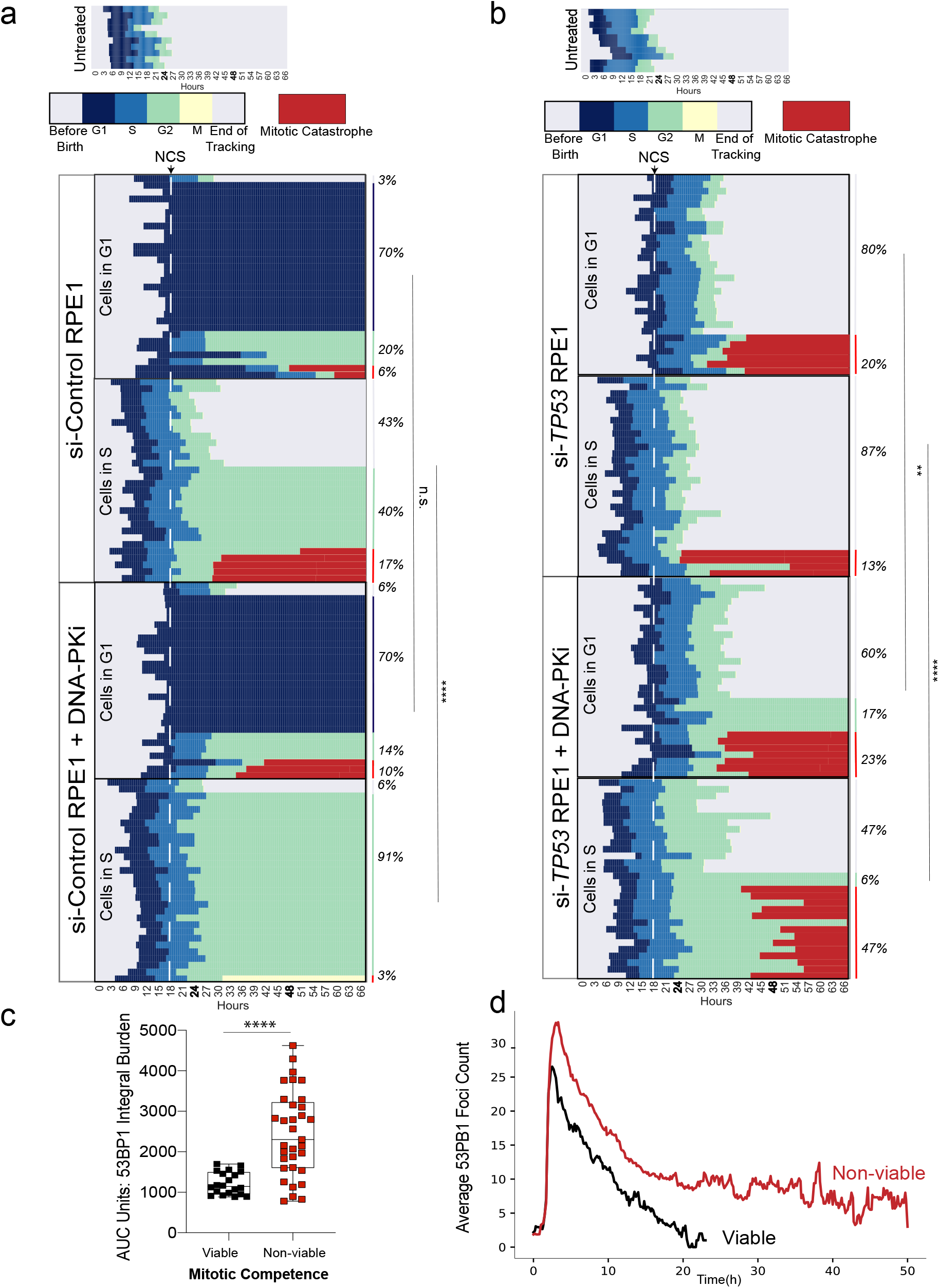
Inhibition of DNA-PK induces catastrophic mitoses in p53-deficient cells. **a,** Cell cycle outcome analyses for si-control treated RPE1, dashed white line indicates drug addition, each row is an individual cell (n = 60 cells for NCS and n=60 cells for NCS+DNA-PKi treatment). Colored bars indicate different phases of the cell cycle, legend shown with no treatment control for comparison. Cells with red bars at the end of mitosis indicate terminal cell cycle event (mitotic catastrophe or apoptosis). Event frequency is reported as a percentage on the right. Cells exposed in G1 vs. S cells are treated as separate cohorts. Fisher’s exact test was performed between −/+ DNA-PKi cohorts using 2 outcome groups (viable, vs. non-viable (arrested cells + terminal outcomes). \*\*\*\**p*<0.0001, n.s. =non-significant **b,** Cell cycle outcome analyses for si-*TP53* treated RPE1, dashed line indicates drug addition, each row is an individual cell (n = 60 cells for NCS and n=60 cells for NCS+DNA-PKi treatment). **c,** AUC analysis of 53BP1 damage burden in viable vs. non-viable p53-deficient cells that were treated with NCS and DNA-PKi. Statistical significance was calculated using a Mann-Whitney test comparing ranks. \*\*\*\**p*<0.0001 **d,** Dynamics of 53BP1 foci burden p53-deficient RPE1 segregated by mitotic viability. The red line corresponds to mean 53BP1 foci burden for all p53-deficient cells treated with NCS and DNA-PKi that undergo catastrophic mitoses, black line indicates mean foci value for p53-deficient cells with NCS and DNA-PKi treatment that are viable post mitosis, (n = 20 viable cells and n = 33 non-viable cells).

Despite the significant increase in mitotic catastrophe induced by combined treatment with DNA-PKi and NCS, 47% of p53-deficient cells exhibit intrinsic resistance to therapy with retained proliferative viability (Fig. 4b). We hypothesized that levels of unrepaired DNA damage may be determinants of viable (*i.e.*, resistant) versus non-viable (*i.e.*, sensitive) cell fates. To evaluate this hypothesis, we quantified integral DNA damage burden in p53-deficient RPE1 with viable versus non-viable mitotic outcomes (Fig. 4c and Supplementary Fig. 4c). The mean integral DNA damage burden was approximately 2-fold higher in non-viable cells, relative to cells that viably completed mitosis (p<0.0001). Further analysis revealed that integral DNA damage burden in S phase was most highly associated with cell viability after drug treatment (Supplementary Fig. 4c). In addition, we traced the average 53BP1 foci burden over time for these two cohorts (Fig. 4d). Our results indicate that cells with non-viable mitotic outcomes have an increased peak value of DNA damage after treatment with DNA-PKi and NCS, which remains elevated over time (p<0.0001 at t=20 hrs, Fig. 4d). Conversely, these findings indicate that p53-deficient cells that exhibit intrinsic therapeutic resistance may be utilizing compensatory DSB repair pathways to counteract the effects of NCS and DNA-PKi prior to mitotic entry.

### p53-deficient cells utilize alternative end-joining pathways in the absence of active DNA-PK

Prior studies have demonstrated that cells with NHEJ deficiency exhibit a compensatory increase in alternative end-joining repair mediated by DNA polymerase theta (Pol θ, gene *POLQ*)^4,33,34^. Polymerase theta dependent end joining (TMEJ) of DNA DSBs is characterized by deletions and templated insertions that are flanked by short tracts of sequence identity, or microhomology (MH)^4^. We found that *POLQ* expression was 10-to 20-fold higher in two independent *TP53^−/−^* RPE1 clones, relative to parental *TP53* wild-type cells (Fig. 5a). *POLQ* is also overexpressed in TCGA breast, lung, bladder, colorectal, gastric, glioblastoma, pancreatic, prostate, melanoma, and uterine cancers with *TP53* mutation, relative to their *TP53* wild-type counterparts (Fig. 5b).

**Figure 5.**
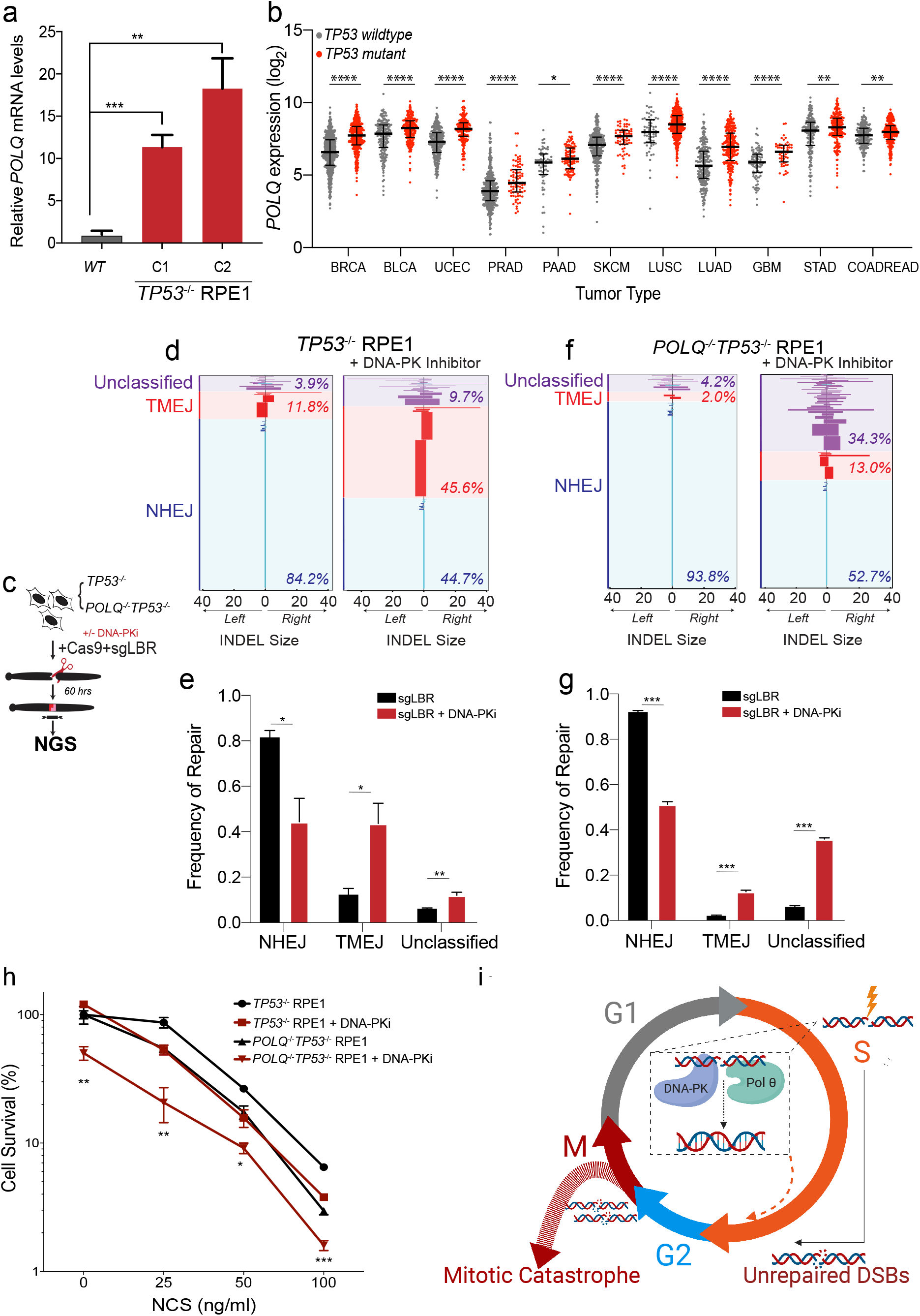
p53-deficient cells utilize alternative end-joining pathways in the absence of active DNA-PK. **a,** RT-qPCR for *POLQ* mRNA levels in 2 *TP53*^−/−^RPE1 clones compared to WT RPE1. Significance was determined using two-tailed t-test. \*\*\*\**p*<0.0001, \*\**p*<0.01. **b,** *POLQ* gene expression depicted as log2 values of *TP53* wild-type vs. mutant cancers across a subset of TCGA tumor types. Tumor labels follow TCGA labeling format. BRCA: breast cancer, BLCA: B-cell lymphoma, UCEC: uterine cancer, PRAD: Prostate cancer, PAAD: pancreatic cancer, SKCM: melanoma, LUSC: lung squamous cell cancer, LUAD: lung adenocarcinoma, GBM: glioblastoma multiforme, STAD: stomach cancer, and COADREAD: colorectal cancer. \*\*\*\**p*<0.0001, \*\*\**p*<0.001, \*\**p*<0.01, *p,0.05, as calculated by one-way ANOVA. **c,** Schematic depicting chromosomal break repair assay. *TP53*^−/−^, and *POLQ^−/−^TP53*^−/−^ RPE1 are segregated into 2 cohorts (+/− 3 uM DNA-PKi). Cells are electroporated using Cas9-RNP-sgRNA-*LBR* and evaluated by next generation sequencing for break repair products at target locus. **d,** Horizontal bar chart representation of individual break repair products at *LBR* locus in *TP53*^−/−^ RPE1 by NGS. Position 0 denotes *LBR* locus cut site, with left and right positions denoting final INDEL size and orientation. Results are reported as average with SEM of n=3 independent biological replicates. **e,** Histogram of overall frequency of repair of NHEJ, TMEJ, and Unclassified products in *TP53*^−/−^ RPE1 with or without DNA-PKi treatment. **f,** Horizontal bar chart representation of individual break repair products at *LBR* locus in *POLQ*^−/−^*TP53*^−/−^ RPE1 by NGS. Position 0 denotes *LBR* locus cut site, with left and right positions denoting final INDEL size and orientation. Results are reported as average with SEM of n=3 independent biological replicates. **g,** Histogram of overall frequency of repair of NHEJ, TMEJ, and Unclassified products in *TP53*^−/−^ RPE1 with or without DNA-PKi treatment. **h,** Colony forming efficiency assay evaluating *TP53*^−/−^ and *POLQ^−/−^TP53*^−/−^ RPE1 after treatment with NCS (at 25 ng/ml, 50 ng/ml, and 100 ng/ml) with or without .5 uM DNA-PKi, data shown are mean +/− SEM (n= 3). Statistical significance assessed with student’s two-tail test. \*\*\**p*<0.001, \*\**p*<0.01, * p<0.05. in comparison to the survival curve of *TP53^−/−^* + DNA-PKi. **i,** Graphical summary

To assess whether hyperactive TMEJ contributes to therapeutic resistance of *TP53^−/−^* RPE1 cells to NCS and DNA-PKi, we sought to inhibit Pol θ. As pharmacological inhibitors of Pol θ are not yet commercially available, we created a double knockout *POLQ^−/−^TP53^−/−^* RPE1 line (Supplementary Fig. 5a). Bi-allelic frameshift mutations in *POLQ* were confirmed by Sanger sequencing and functional deficiency was established using an extrachromosomal TMEJ repair assay (Supplementary Fig. 5b-d)^4^.

To directly assess whether TMEJ repair is increased after DNA-PKi treatment, we analyzed chromosomal break repair patterns at a site-specific DSB in p53-deficient RPE1 cells. Cells were transfected with Cas9 ribonucleoprotein (RNP) complexes that target the *LBR* locus, with or without DNA-PKi^35^. Genomic DNA was harvested 60 hours later and analyzed for break repair patterns using next generation sequencing (NGS) (Fig. 5c). Target amplification and TIDE analyses confirmed high rates of target site cleavage in all samples transfected with a full complement of Cas9-RNP (Supplementary Fig. 5e,f)^36^. We applied a bioinformatic algorithm (ScarMapper, see methods) to characterize the spectrum of repair products with at least 0.1% prevalence, classified according to the size of left deletion (LD), right deletion (RD), insertion (Ins), and microhomology (MH) (ScarMapper Methods). Indels <5bp were categorized as NHEJ, with the predominant repair product being a +A 1bp insertion^35^. TMEJ was defined as repair products whose frequency was diminished by at least 2-fold in *POLQ^−/−^* cells. All other repair products were categorized as “Unclassified.” DNA-PK inhibition in *TP53^−/−^* RPE1 cells results in a substantial reduction in NHEJ repair, with a compensatory increase in TMEJ to nearly 45% of all DSB repair (Fig. 5d,e). In contrast, DNA-PK inhibition in *POLQ^−/−^TP53^−/−^* RPE1 cells did not result in a substantial increase TMEJ signature repair (Fig. 5f,g). However, a higher proportion of Unclassified repair products were detected (Fig. 5f,g). A limitation of NGS analysis of DSB break repair is that non-amplifiable target loci are not measured. Thus, we used digital PCR to quantify the *LBR* locus detection rate, relative to a control locus, upon inhibition of DNA-PK and/or Pol θ (Supplementary Fig. 5g,h). *LBR* locus detection rates were most reduced upon inhibition of DNA-PK and Pol θ, consistent with overall inhibition of DSB repair (Supplementary Fig. 5h). These observations confirm an essential role for TMEJ in compensatory repair of chromosomal DSBs upon pharmacologic inhibition of DNA-PK.

To determine the impact of *POLQ* inhibition on cellular viability, we performed clonogenic survival assays in the parental *TP53^−/−^* and *POLQ^−/−^TP53^−/−^* RPE1 lines treated with NCS with or without DNA-PKi. Genetic deficiency in *POLQ* resulted in significantly reduced viability after NCS treatment, particularly in combination with DNA-PKi (Fig. 5h). *TP53*^−/−^RPE1 cells with inhibition of both TMEJ and NHEJ repair pathways had comparable clonogenic survival to p53-proficient RPE1 cells (see Supplementary Fig. 1e). Collectively, these findings indicate that hyperactive end joining repair via NHEJ and TMEJ mediate resistance to DNA damaging therapy induced by p53 deficiency (Fig. 5i).

## Discussion

These results recognize enhanced DNA end joining repair capacity as a novel component of therapeutic resistance induced by p53 deficiency, and that loss of functional p53 alone is sufficient to increase hyperactive repair. Our findings indicate that DSB end joining hyperactivity is particularly relevant for suppressing S phase DNA damage burden, which we find is a key determinant of mitotic catastrophe (Fig. 5i). Although NHEJ Is conventionally considered to be most critical for repair in G1, we observed a relatively greater impact of DNA-PK inhibition on the fate of S phase cells after treatment with a radiomimetic. There are several potential explanations for this unanticipated observation. First, recent findings suggest that DNA-PK may be dispensable for synapsis formation during NHEJ^37^. Accordingly, repair of “simple” DSBs in G1 phase may have a reduced reliance on DNA-PK, whereas repair of more “complex” DSBs in S phase may require DNA-PK, possibly in partnership with the nuclease Artemis^38–40^. Second, it is possible that DNA-PK inhibition may be more impactful in S phase due to trapping of Ku proteins at DSBs, which inhibits the activation of homologous recombination pathways^41^. Third, DNA-PK may be particularly important in early S phase, when sister chromatids are not broadly present. Notably, we observed a prominent peak of unrepaired DSBs just as p53-deficient cells transitioned from G1 to S phase. Our observation that DSB end joining hyperactivity in p53-deficient cells is highly sensitive to DNA-PK inhibition warrants further mechanistic investigation. Recently, CYREN (cell cycle regulator of NHEJ) has been proposed to be a cell-cycle phase specific inhibitor of the Ku70/80 heterodimer that is critical for restricting NHEJ to G1^42^. It is therefore possible that p53-deficiency may transcriptionally reprogram cell cycle-inhibitors of NHEJ to enable hyperactive repair, though that is beyond the scope of this study.

Regulatory mechanisms that confer TMEJ hyperactivity in cancer are not well understood, although transcriptional overexpression of *POLQ* has also been observed in breast and ovarian cancers with *BRCA1*/*BRCA2* deficiency or mutations in other genes that confer Pol γ synthetic lethality^33,43^. Recent work investigating integrated pathway analysis of *TP53* deficiency noted *POLQ* to be frequently overexpressed in *TP53* pathway deficient cancers^1^. Our findings, in an isogenic p53-deficient cell line model, indicate that this relationship may be causal. The mechanism for p53-dependent suppression of *POLQ* expression remains to be elucidated, and may entail the regulation of non-coding RNAs^44^. The use of TMEJ can also be explained by the potential creation of more complex DSBs upon NHEJ suppression that serve as poor substrates for homologous recombination (HR). Indeed, the molecular mechanisms of NHEJ and TMEJ hyperactivity induced by p53 deficiency warrant further investigation.

Radiotherapy and other forms of DNA damaging therapy are employed in the vast majority of cancer patients^45^. Resistance to DNA damaging therapy may thus explain the adverse clinical outcomes associated with *TP53* mutations in many different cancer types^1^. Our study supports the investigation of DNA-PK inhibitors administered in combination with DNA damaging therapy (including radiotherapy) in patients with p53-deficient cancers. Additionally, as inhibitors of Pol θ are currently in development^46^, our study suggests that combined inhibition of both DNA-PK and Pol θ represents a promising strategy to reverse the therapeutic DNA damage resistance in p53-deficient cancers.

## Materials and Methods

### Key Resources Table

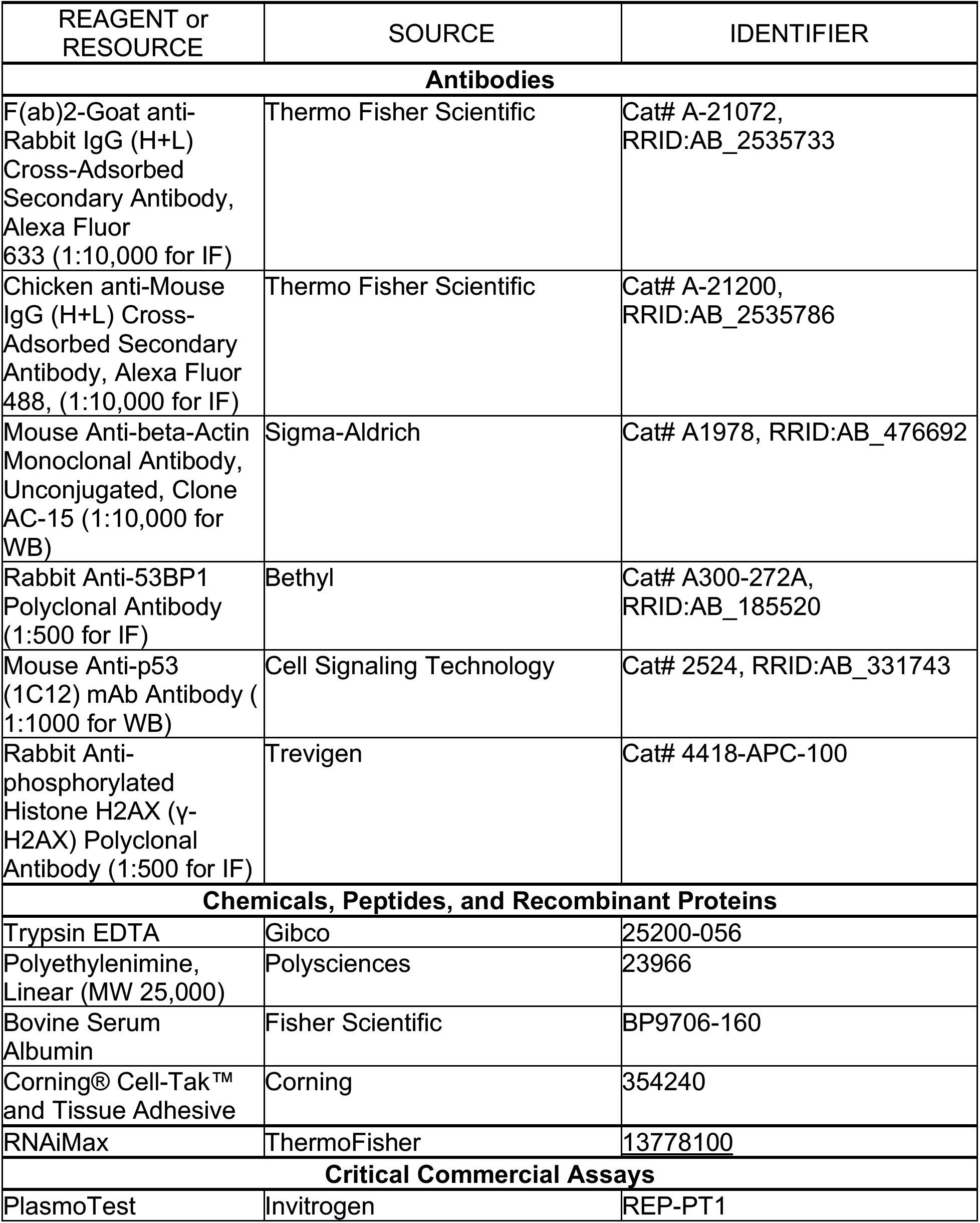
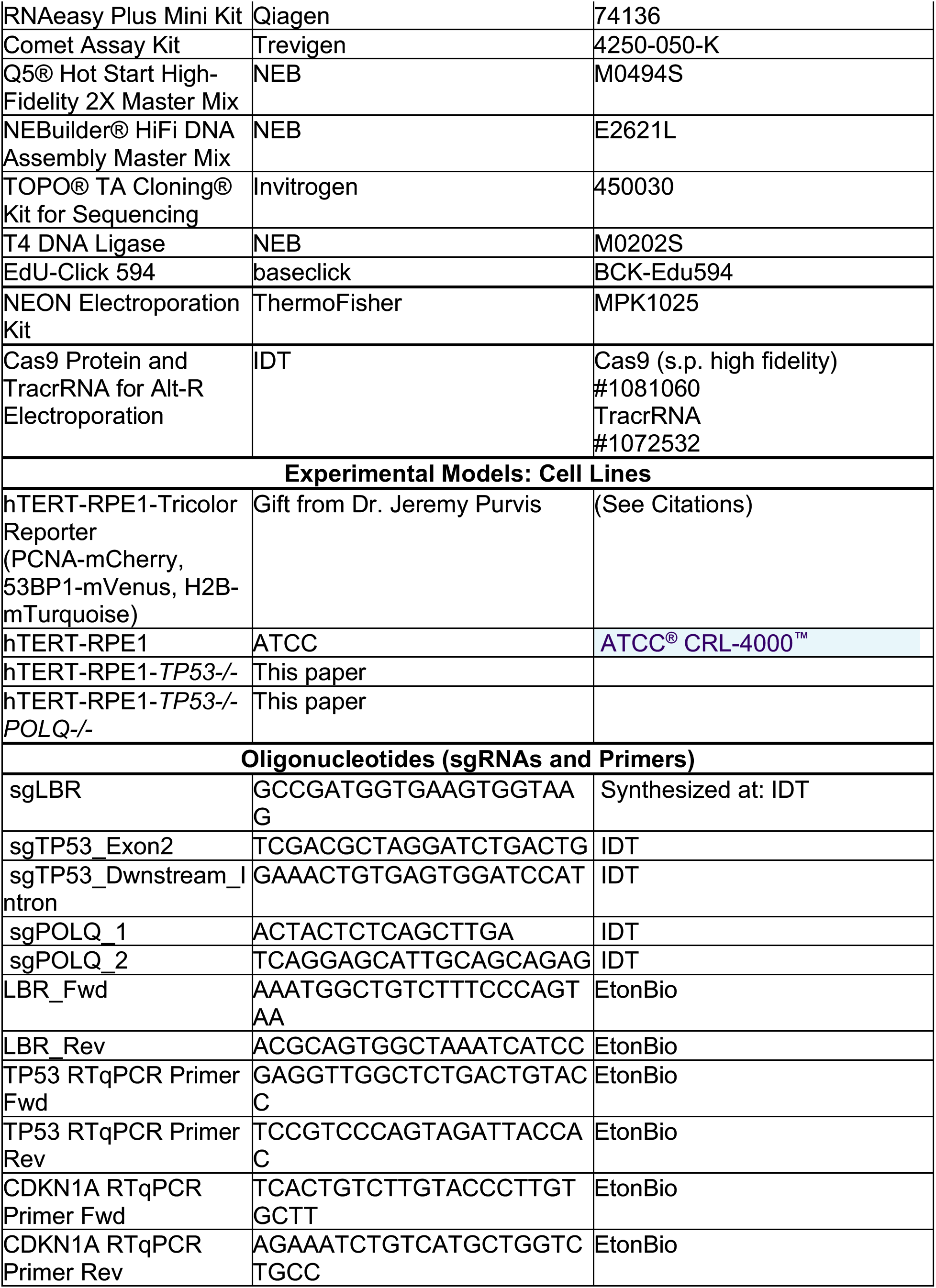
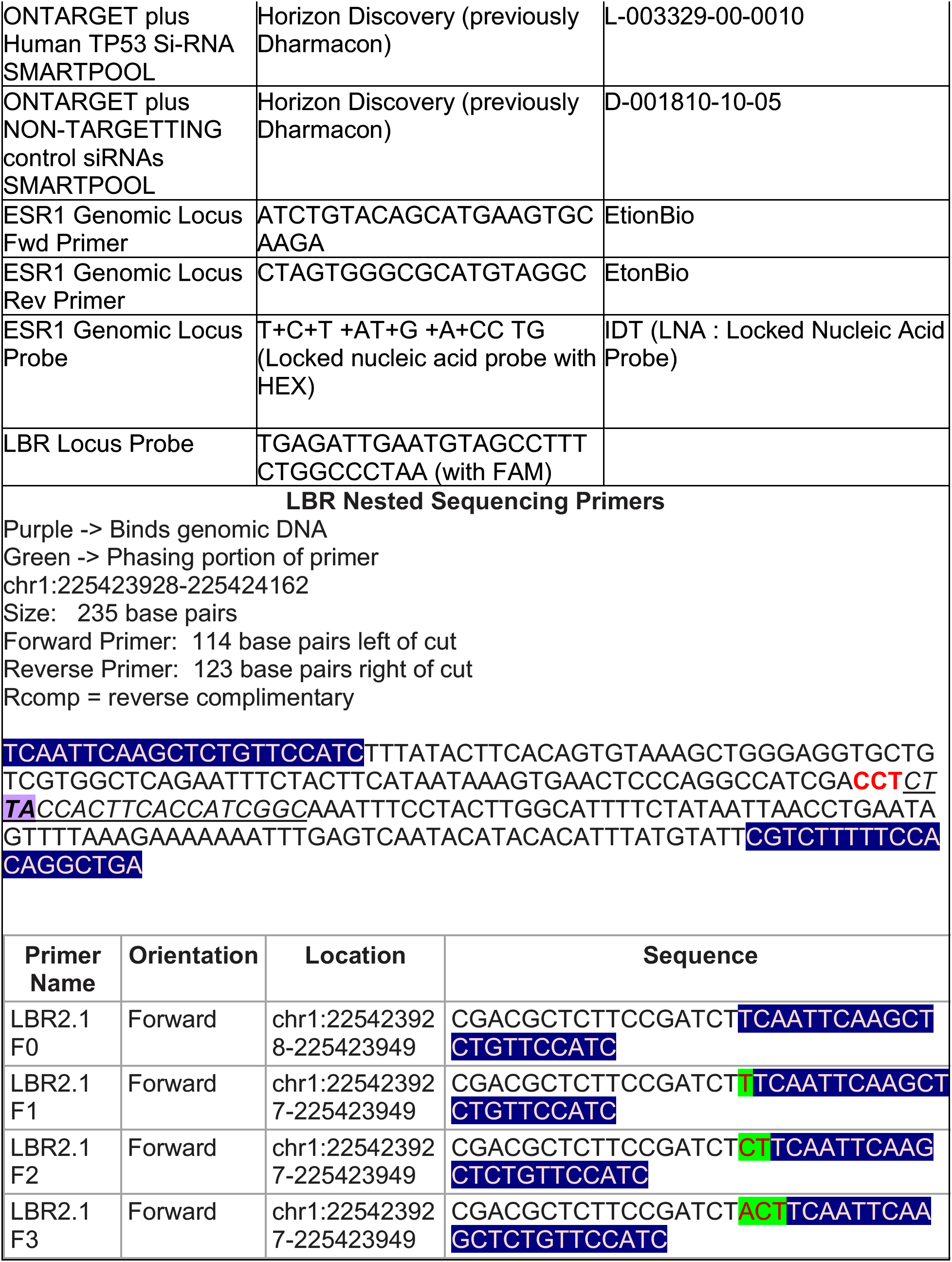
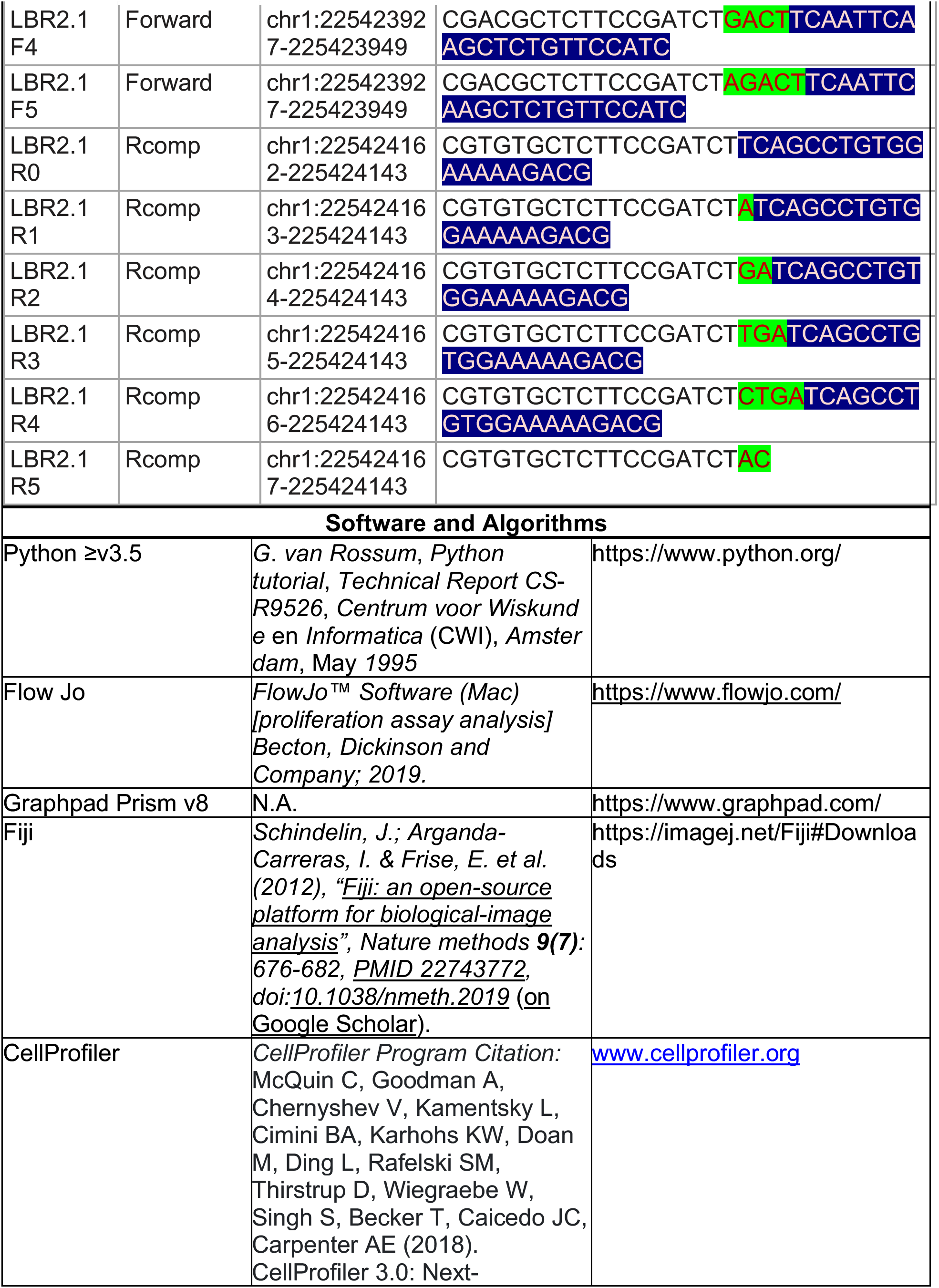
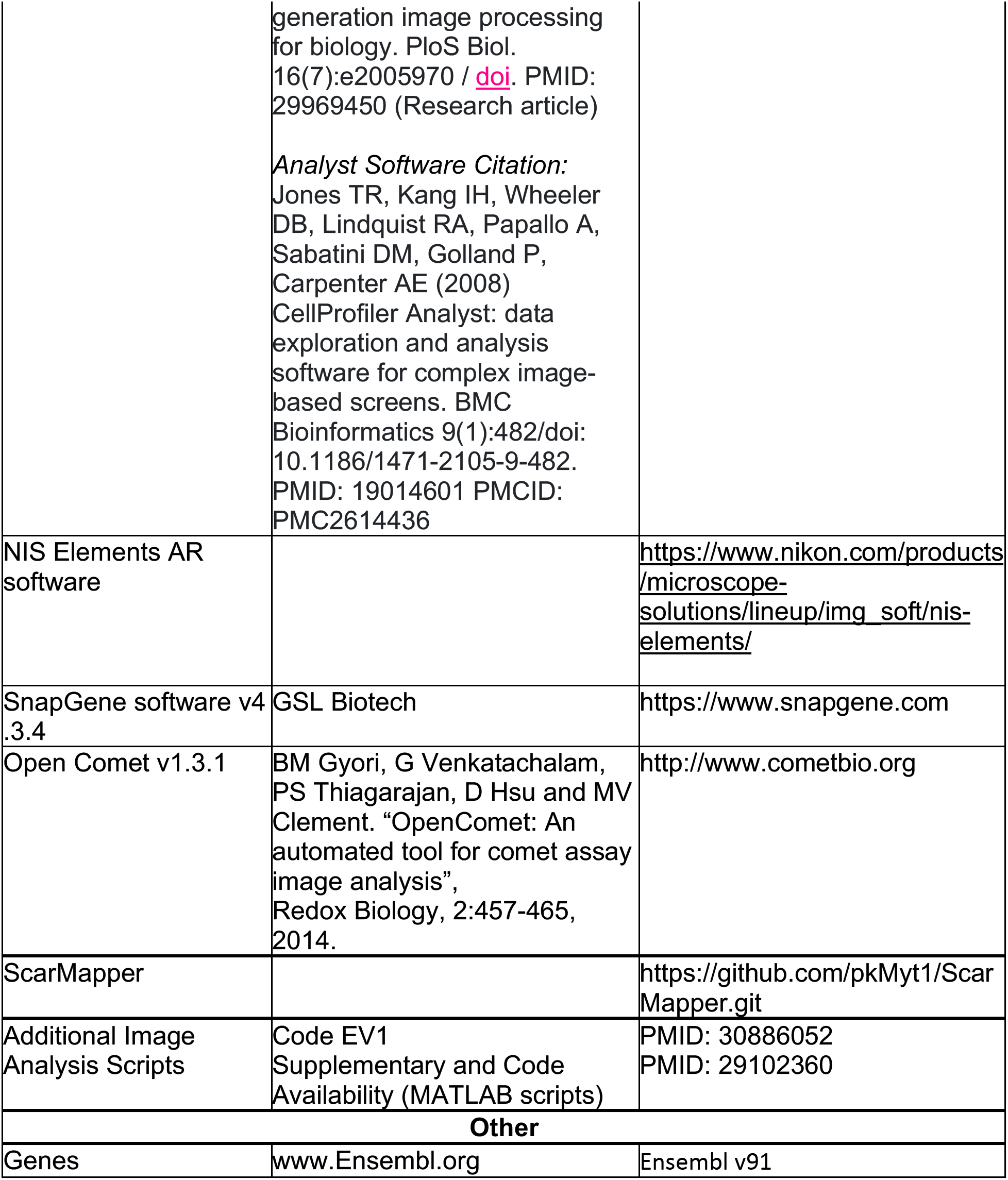

#### Cell Culture

*WT (p53^+/+^), WT Fusion-Reporter (p53^+/+,^PCNA-mCherry, 53BP1-mVenus), (p53^−/−^,* and *p53^−/−^Polq^−/−^*cells are hTERT immortalized RPEs. Cells were maintained in Dulbecco’s modified Eagle’s medium (DMEM), with 10% Fetal Bovine Serum (Hyclone FBS) and 2mM L-glutamine (ThermoFisher).All cells were maintained at 37 C in an atmosphere of 5% CO2. Cells were routinely tested for mycoplasma contamination using PlasmoTest (Invivogen).

#### Establishment of Stable Cell Lines

For the TP53 and Polq mutant cell lines, we used the Alt-R-CRISPR-Cas9 system (IDT). We performed Neon transfection (Invitrogen) and followed the manufacturer’s protocol with Alt-R HiFi Cas9 nuclease, crRNA and tracrRNA purchased from IDT. crRNA was designed using MIT CRISPR (http://crispr.mit.edu) to target Exon 2 of the TP53 gene for the p53 mutant cell line and the polymerase domain of the Polq gene for the Polq mutant cell line. Forty-eight hours after transfection, cells were seeded for single clone selection. For the p53 gene editing experiment, a homologous template with a stop codon and SCA-I site was provided for selection of gene edited cells. Restriction Enzyme screening, PCR screening, and Sanger sequencing confirmed gene targeting, post which we performed functional tests.

#### Immunofluorescence

Cells were fixed with 3% Paraformaldehyde for 15 min at RT, followed by permeabilization with 0.25% TritonX-100 in PBS. Cells were subsequently processed for immunostaining experiments using the antibodies listed below. Nuclei were visualized by staining with DAPI. The primary antibodies used were: γH2AX (1:500, Trevigen, 4418-APC-100), and 53BP1 (1:500 for immunofluorescence, Bethyl, A300-272A). The secondary antibodies were: FITC Goat Anti-Mouse IgG (H+L) (1:500, Jackson ImmunoResearch, 115-095-003) and FITC Goat Anti Rabbit IgG (H+L) (1:500, Jackson ImmunoResearch, 111-095-144). Images were acquired using the GE IN CELL 2200 high through-put imaging system at 40x magnification.

#### siRNA Treatment

WT Fusion-Reporter RPE cells were passaged twice after −80 thaw and plated on 12 well plates at a density of 100,000 cells/ well for siRNA treatment. Twenty-four hours post plating, cells were exposed to 10nM / well sip53 (SMART pool from Dharmacon), and siControl (Non-targetting SMART pool from Dharmacon), in OPTIMEM with RNA-iMAX (ThermoFisher) as a transfection reagent. As a no-treatment control, cells were exposed to RNA-iMAX and OPTIMEM without siRNA. 48 hours post transfection, cells were transferred onto 12 well Cell-Tak coated glass plates (Cellvis), at a concentration of 50,000 cells/well for imaging. Prior to imaging and at the end of imaging, samples were taken for RT-qPCR analysis of p53 mRNA to confirm si-RNA knockdown.

#### Mixed Competition Assay-Flow Cytometry

mCherry labelled and unlabeled hTERT-RPE1 cell lines were plated on 96 well plates at a 50:50 ratio, and irradiated 2 hrs post plating at 0, 2, 4, or 6 Gy, and left to grow. At indicated timepoints cells were harvested by trypsinizing and quenching with PBS with 5% BSA. Cells were fixed with 2% PFA and subsequently transferred to V-bottom plates (ThermoFisher, 249570). Cells were quantified by flow cytometry using the Intellicyt iQue at a volume of 100 ul / sample, collecting all events per well. For each condition, 6 biological replicates.

#### Time-Lapse Imaging Microscopy

Cells stably expressing Proliferating Cell Nuclear Antigen (PCNA)-mCherry and Tumor Suppressor p53 Binding Protein 1 (53BP1) – mVenus were treated with si-RNA for 48 hours prior to imaging. PCNA-mCherry and 53BP1-mVenus fusion reporter is a gift from Dr. Jeremy Purvis and Hui Chao Xiao. Cells were plated on Cell-Tak (Corning) coated glass-bottom 12-well plates (Cellvis) with Phenol-free DMEM (Invitrogen) supplemented with 10% FBS, and L-glutamine. Twenty-four hours post plating, cells were image captured every 10min for 72h in the mCherry and mVenus fluorescence channels. 18 hours into imaging, DNA PKi was added at a concentration of 0.5 uM / well, and/or NCS at a concentration of 100ng/ mL/ well. We commenced imaging every 10 minutes in both channels for another 48 hours. Fluorescence images were obtained using a Nikon Ti Eclipse inverted microscope with a 40x objective and Nikon Perfect Focus (PFS) system to maintain focus during acquisition period. Cells were maintained at constant temperature (37°C) and atmosphere (5% CO_2_). Nikon, NIS Elements AR software was utilized for image acquisition. Image analysis was performed on ImageJ – Fiji and Cell Profiler.

#### Colony Forming Assays

Cells lines used in the assay are indicated in the figures. Cells were treated with NCS and/ or DNA-PKi for twenty-four hours, after which we performed a media change. For colony formation experiments with ionizing radiation, cells were plated for IR treatment with or without DNA-PKi, and inhibitor treatment was washed off after twenty-four hours. Cells were subsequently incubated for 10-12 days at 37°C to allow colony formation. Colonies were stained by Coomassie blue and counted.

#### DNA Repair Assay

Cell lines used in the assay are indicated in the figure. 5 × 10^5^ cells were transfected with sgLBR2 and TracrRNA complexed Cas9 protein at final concentrations of sgRNA:tracrRNA duplex: 22 pmol and Cas9: 18 pmol per reaction, with Neon transfection kit (Invitrogen) using 2 1350V, 30ms pulses in a 10μL chamber. 60 hours post transfection, cells were harvested for genomic DNA extraction (Nucleospin). Part of the gDNA was utilized for Sanger Sequencing and TIDE analysis post amplification of the genomic LBR2 locus. Remaining gDNA was amplified using NGS nested sequencing primers and sent for sequencing and/ or Digital PCR.

#### Digital PCR

Primers and 5’ hydrolysis probes were designed to specifically detect the copies of *LBR* locus. *ESR1* locus was used as genomic control. Each reaction assay contained 10 μLof 2x dPCR Supermix for Probes (No dUTP), 0.9 μmol/L of respective primers, 0.25 μmol/L of respective probes, and 10 ng of DNA in a final volume of 20 μL. Droplets were generated using automated droplet generator (Bio-Rad catalog #186-4101) following manufacturer’s protocol. PCR parameters for *LBR* locus were 10 sec at 95 °C, then 40 cycles of 94 °C for 30 sec, 60 °C for 30 sec, and 72°C for 2 min followed by 98°C for 10 min with a ramping of 2 °C/sec at all steps. The PCR cycling parameters for *ESR1* genomic locus were 10 sec at 95 °C, then 40 cycles of 94 °C for 30 sec and 60 °C for 1 min followed by 98°C for 10 min with a ramping of 2 °C/sec at all steps. After PCR amplification, droplet reader (Bio-Rad QX200™ Droplet Reader Catalog #1864003) was used to measure the end-point fluorescence signal in droplets as per the manufacturer’s protocol. The recorded data was subsequently analyzed with QuantaSoft software version 1.7.4.0917 (Bio-Rad). Each Taqman probe was evaluated for sensitivity and specificity.

#### DNA Repair Assay

Cell lines used in the assay are indicated in the figure. 5 ×10^5^ cells were transfected with sgLBR2 and TracrRNA complexed Cas9 protein at final concentrations of sgRNA:tracrRNA duplex: 22 pmol and Cas9: 18 pmol per reaction, with Neon transfection kit (Invitrogen) using 2 1350V, 30ms pulses in a 10μL chamber. 60 hours post transfection, cells were harvested for genomic DNA extraction (Nucleospin). Part of the gDNA was utilized for Sanger Sequencing and TIDE analysis post amplification of the genomic LBR2 locus. For analysis of INDELs, 100 ng of gDNA was amplified using phased primers. These libraries were indexed with the Illumina unique dual combinatorial indices. Following pooling, 2 × 150 cycle sequencing was done on an Illumina iSeq™. INDELs were identified by comparing the target reference sequence to the resulting sequence reads in the FASTQ files via a 10-nucleotide sliding window using the ScarMapper program.

## Acknowledgements

We thank Kasia Kedziora, Samuel Wolff, and Juan Carvajal-Garcia for data acquisition and technical assistance. We are grateful to the Gupta and Purvis Lab members for helpful discussions. UNC Core labs (Microscopy Services Laboratory, Hooker Imaging Core, and Flow Cytometry Core Facility) used in this study are supported in part by P30 CA016086 Cancer Center Core Support Grant to the UNC Lineberger Comprehensive Cancer Center. Funding support was provided by the NCI/NIH (CA222092), Dept of Defense (W81XWH-18-1-0047), and the University Cancer Research Fund. G.P.G. holds a Career Award for Medical Scientists from the Burroughs Wellcome Fund. R.J.K. is supported by the Cancer Cell Biology T32 Training Program (2T32CA071341-21) and the UNC Medical Scientist Training Program (MSTP).

## Author contributions

R.J.K and G.P.G. designed and conceived experiments. G.P.G. supervised the study. R.J.K and H.C.X performed the live-cell imaging experiments. H.C.X. and J.E.P. provided critical reagents and image analysis guidance. R.J.K performed and implemented computational analyses for image processing. R.J.K. performed all additional experiments and data analyses with statistical review. V.R.R., A.R.S., S.J.S., W.F., A-S.W., and S.K. provided technical assistance on imaging acquisition, colony forming assays, and digital PCR. D.A.S. developed break site sequencing analysis platforms. R.J.K. and G.P.G. wrote the manuscript, with contributions from all authors. All authors read and accepted the manuscript.

---

**Supplementary Figure 1.**
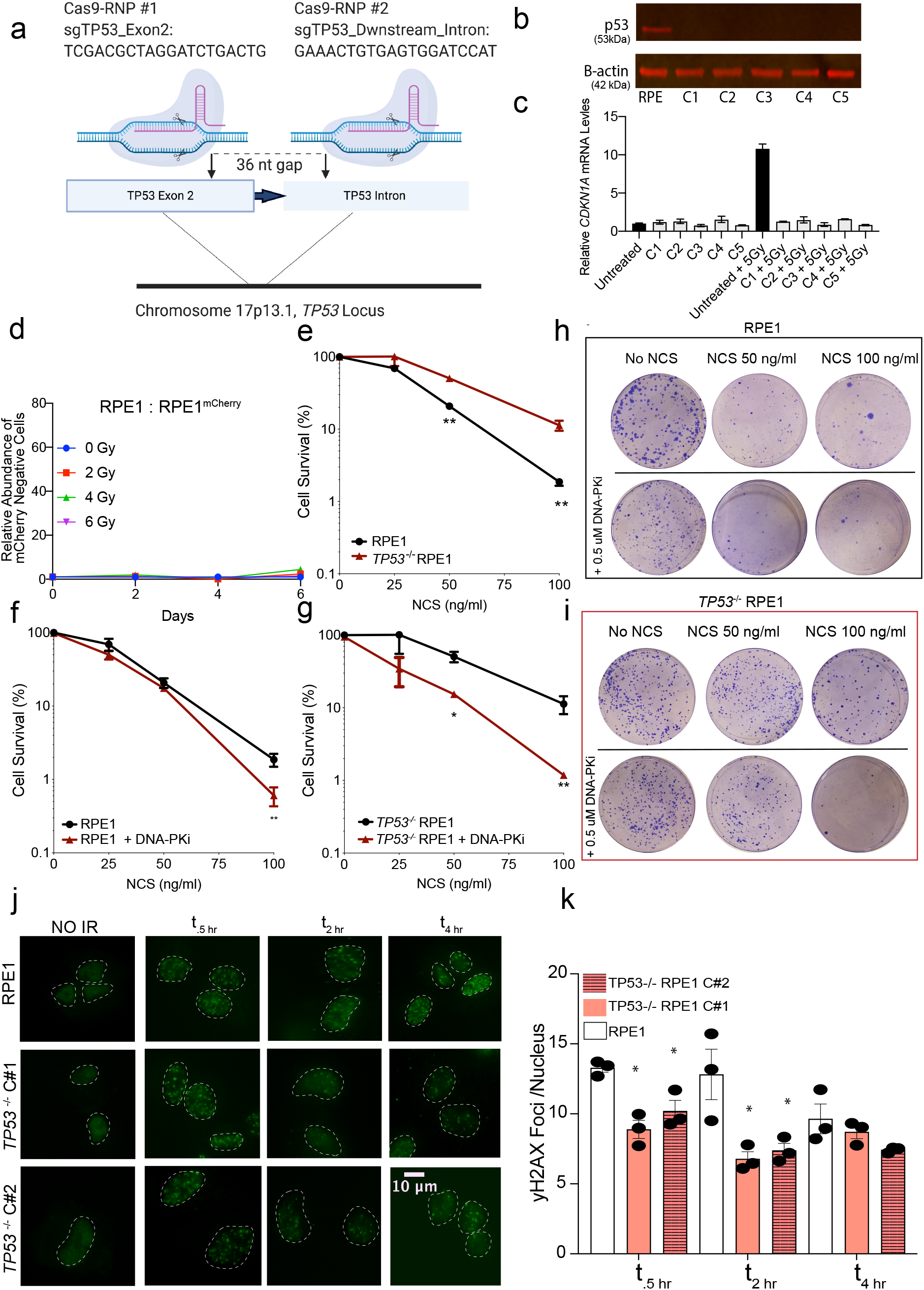
**a,** Schematic of CRISPR target locus in human *TP53* gene. Two sgRNAs were designed to target sites in the terminal region of exon 2 (which encodes the p53 transactivating domain) and a site in the downstream intron with a 36 nucleotide (nt) gap. sgRNAs were complexed with Cas9 in the RNP system and electroporated into RPE1 cells. **b,** Western Blot of 5 selected single-cell clones that were profiled for p53 protein. **c,** Functional assay evaluating p53-dependent *CDKN1A* transcriptional responses to treatment of 5Gy IR. RNA from cells exposed to IR were harvested 6 hrs post treatment. **d,** Relative abundance of unlabeled RPE1^unlabelled^ over RPE1^mCherry^ measured by Intellicyte high-throughput cytometry +/− SEM (n=6) is shown, normalized to the untreated (0Gy) cohort at each time point. **e,** Clonogenic survival assays performed in RPE1 vs *TP53^−/−^* RPE1 cells exposed to NCS. **f,** Clonogenic survival assays of RPE1 treated NCS +/− 0.5 uM DNA-PKi. Reported values are mean of n = 3 replicates, and survival fraction was calculated by first calculating plating efficiency and normalizing it to the untreated samples. **g,** *TP53^−/−^* RPE1 cells were treated with NCS +/− 0.5 uM DNA-PKi. **h,i,** Representative colony forming plates for e and f at NCS doses of 0, 50 and 100 ng/ mL +/− 0.5 uM DNA-PKi. Cell numbers for each conditions plated are the following: UT (500), NCS 50 ng/ ml (2000), NCS 100 ng/ ml (6000). **j,** Representative immunofluorescence images of yH2AX foci in cells with indicated genotypes untreated (no IR) or treated with IR (5Gy) and collected at .5, 2, and 4 h after irradiation. **k,** Quantification of yH2AX foci. Data shown are mean (n= 50 cells per treatment condition) +/− SEM (n=3), and are consistent across two independent biological replicates. \**p*<0.05 by two-tailed Student’s t-test.

**Supplementary Figure 2.**
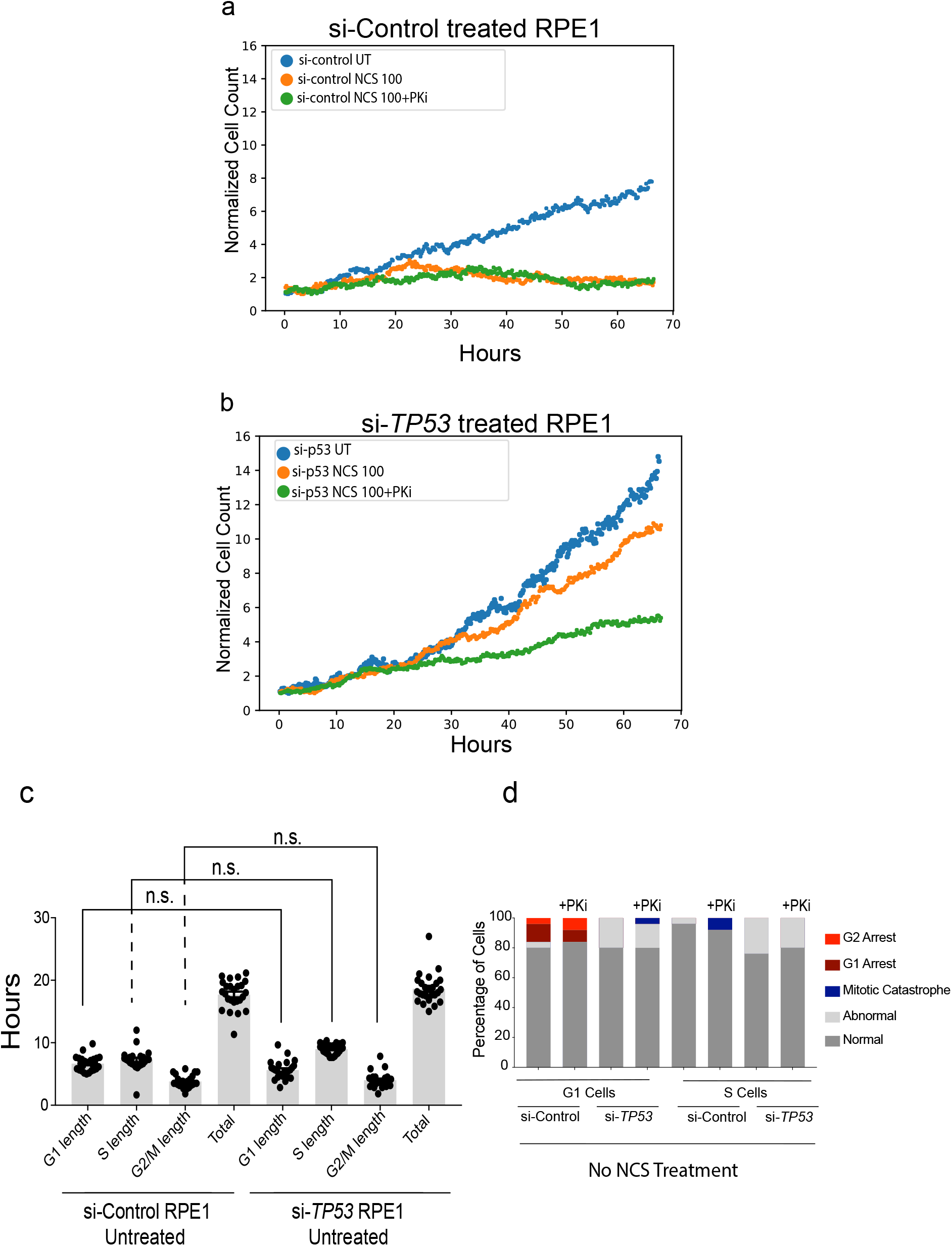
**a,** Quantification of cell proliferation from live-cell imaging experiments for si-Control treated RPE1. Cell counts were normalized to cell numbers at start of imaging. Here we show one representative imaging beacon for each treatment condition (untreated, NCS 100 ng/ ml at 18 hours, and NCS 100 ng/ml + 0.5 uM DNA-PKi at 18 hours). **b,** Cell proliferation counts for si-*TP53* treated RPE1 over live-cell imaging. **c,** Analysis of RPE1 with no exposure to NCS (untreated) but received si-Control or si-*TP53* 3 days prior to imaging. No significant differences are seen due to si-treatment alone. **d,** Quantification of baseline mitotic outcomes with no NCS exposure over the course of imaging. DNA-PKi treatment alone with no NCS showed no little to no additional effect on cells (Chi-squared analysis P>0.05 for each condition).

**Please refer to the following file names for the Supplementary Figure 3 Video Files:** Each video is a split screen of the same cell depicted in 2 channels: PCNA (left) and 53BP1 (right)

**3a:** NormalMitosis.mp4

**3b:** TransientG2Delay.mp4

**3c:** G1Arrest.mp4

**Supplementary Figure 3.** **a, Normal Mitosis:** RPE1 Cell cycle representative of normal mitosis, with NCS treatment only. For all cells in this figure both the PCNA and the 53BP1 channels are shown as two individual movies. **b, Transient G2 Delay:** RPE1 cell cycle representative of a transient cell cycle delay in G2 (length of G2 is significantly prolonged in comparison to untreated cells). This cell was treated with NCS and DNA-PKi. **c, G1 Arrest:** RPE1 cell cycle representative of a permanent G1 arrest. This is a p53 proficient cell treated with NCS and DNA-PKi.

**Supplementary Figure 4.**
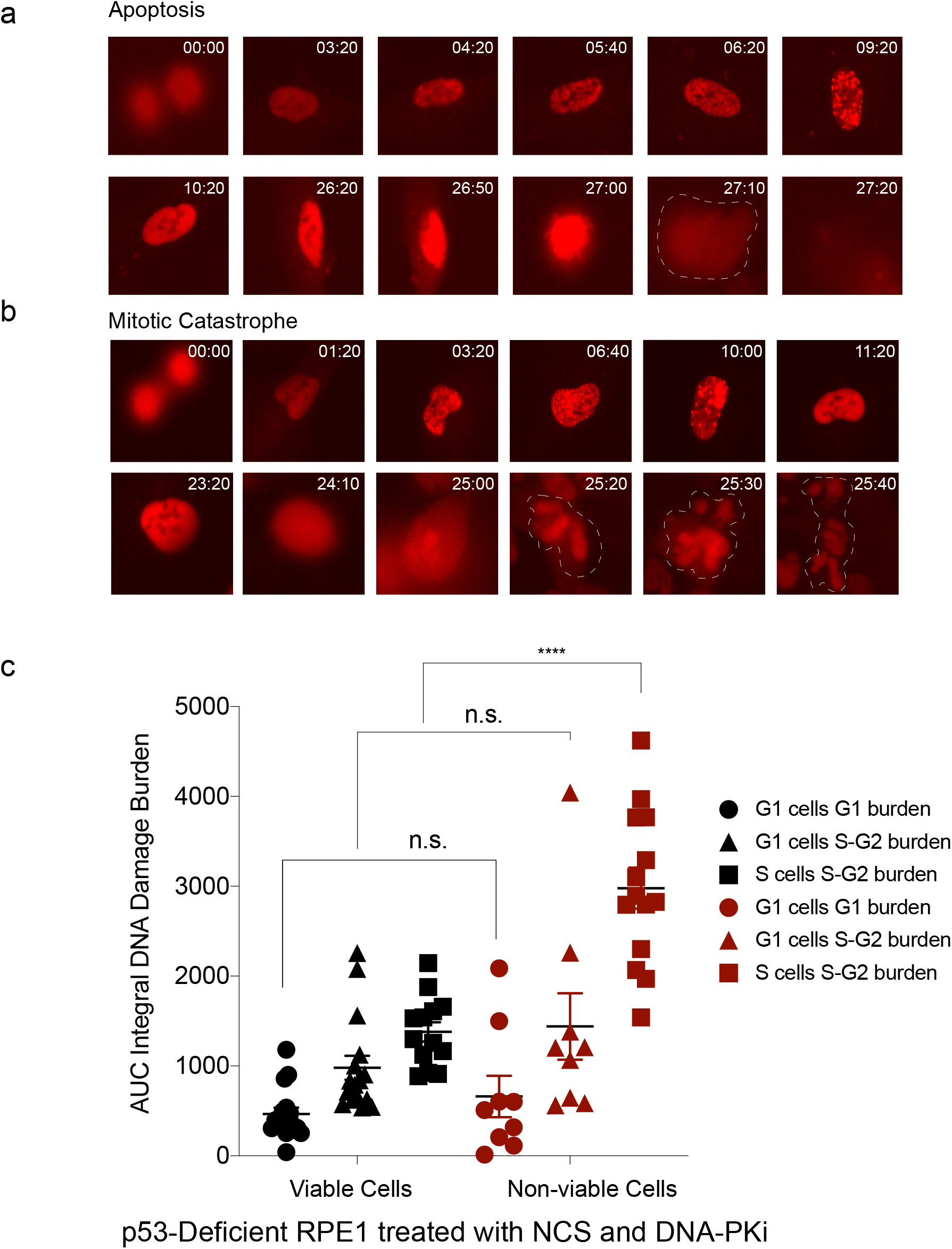
**a,** Time stamped image sequence of apoptotic cell (PCNA channel shown). Cells that experienced nuclear degradation during cell cycle prior to mitosis were categorized as “apoptotic cells.” In this sequence a cell in G2 experiences cell death at 27 hours post birth, with indication of mitotic attempt, with nuclear envelope collapse or presence of any daughter cells. **b,** Time stamped image sequence of cell that experienced mitotic catastrophe (PCNA channel shown). Cell undergoes nuclear envelope collapse (24:10), and attempts mitosis, in subsequent images fragmentation of nucleus is clearly visible with no viable daughter cells present. Cell non-viability during mitosis was defined as mitotic catastrophe. **c,** Integral DNA damage burden for p53-deficient cells treated with NCS (100 ng/ml) and DNA-PKi (.5 uM) are calculated and segregated by viable (black) vs. non-viable outcomes (red). Legend indicates which phase of cell cycle the cells are in during drug exposure, followed by the phase for which the burden is calculated. Ex: G1 cells G1 burden = cells in G1 during drug exposure and total damage burden in G1. Area under the curve (AUC) analysis was performed by plotting 53BP1 foci counts over time for each cell and integrating burden over time. Statistical significance was determined using two-tailed Student’s t-test.

**Supplementary Figure 5.**
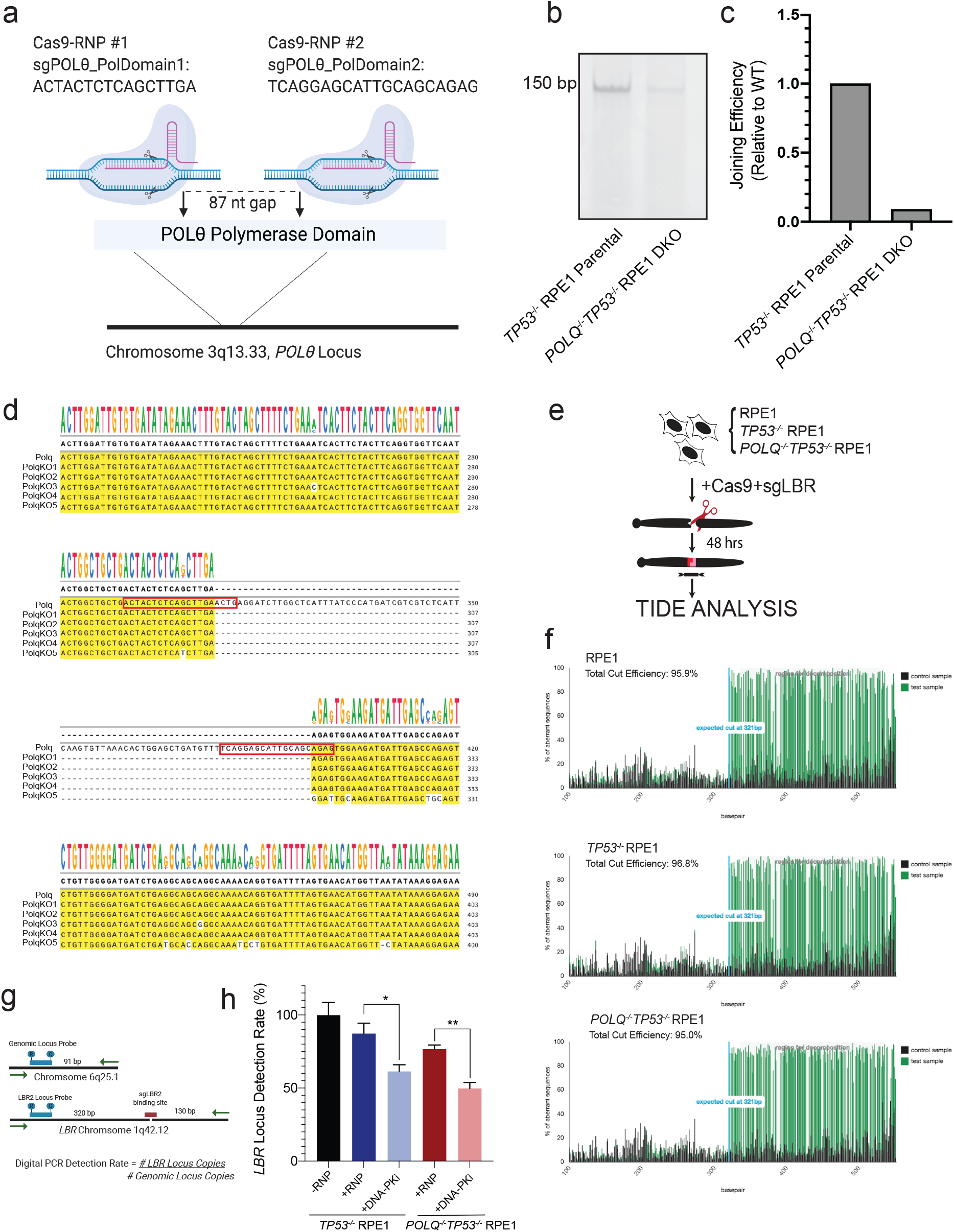
**a,** Schematic of CRISPR target locus in human *POLQ* gene. Two sgRNAs were designed to target sites in the polymerase domain, with an 87 nucleotide (nt) gap. sgRNAs were complexed with Cas9 in RNP system and electroporated into RPE1 cells with a *TP53^−/−^* background to create double knockout cell line. **b,c** POLQ specific substrates were introduced into the *TP53 ^−/−^* vs. *POLQ^−/−^TP53^−/−^* DKO cells to assess repair efficiency. Products were amplified and characterized by electrophoresis and end joining efficiency was normalized to RPE1 with *POLQ* expression. **d,** Sanger sequencing analysis of CRISPR edited locus in *POLQ^−/−^ TP53^−/−^* RPE1 clones. The locus of interest was PCR-amplified and cloned into a TOPO vector for sequencing analyses. Each line of sequence shown was derived from a different TOPO clone and aligned to show differences. The *POLQ^−/−^TP53^−/−^* clone has 87bp deletion resulting in frameshift mutations. Red boxes indicated sgRNAs used for the CRIPSR. **e,** Schematic showing evaluation of NGS samples by TIDE analysis for efficiency of cleavage at *LBR* target site across cell lines. **f,** RPE1, *TP53^−/−^*, and *POLQ^−/−^TP53^−/−^* RPE1 cutting efficiency, all three cell lines have comparable levels of cutting efficiency with sgLBR. **g,** Schematic depicting digital PCR method for assessing LBR detection rate. **h,** Results of digital PCR assay on *TP53 ^−/−^* vs. *POLQ^−/−^TP53^−/−^* cells with −/+RNP and subsequently −/+ DNA-PKi (3uM). Average of 3 independent biological replicates are shown with SEM. Statistical significance was calculated using Multiple-t tests. *p<0.05, **p<0.01.

